## Supplementary Figures for "SFSWAP is a negative regulator of OGT intron detention and global pre-mRNA splicing"

**Fig. S1**

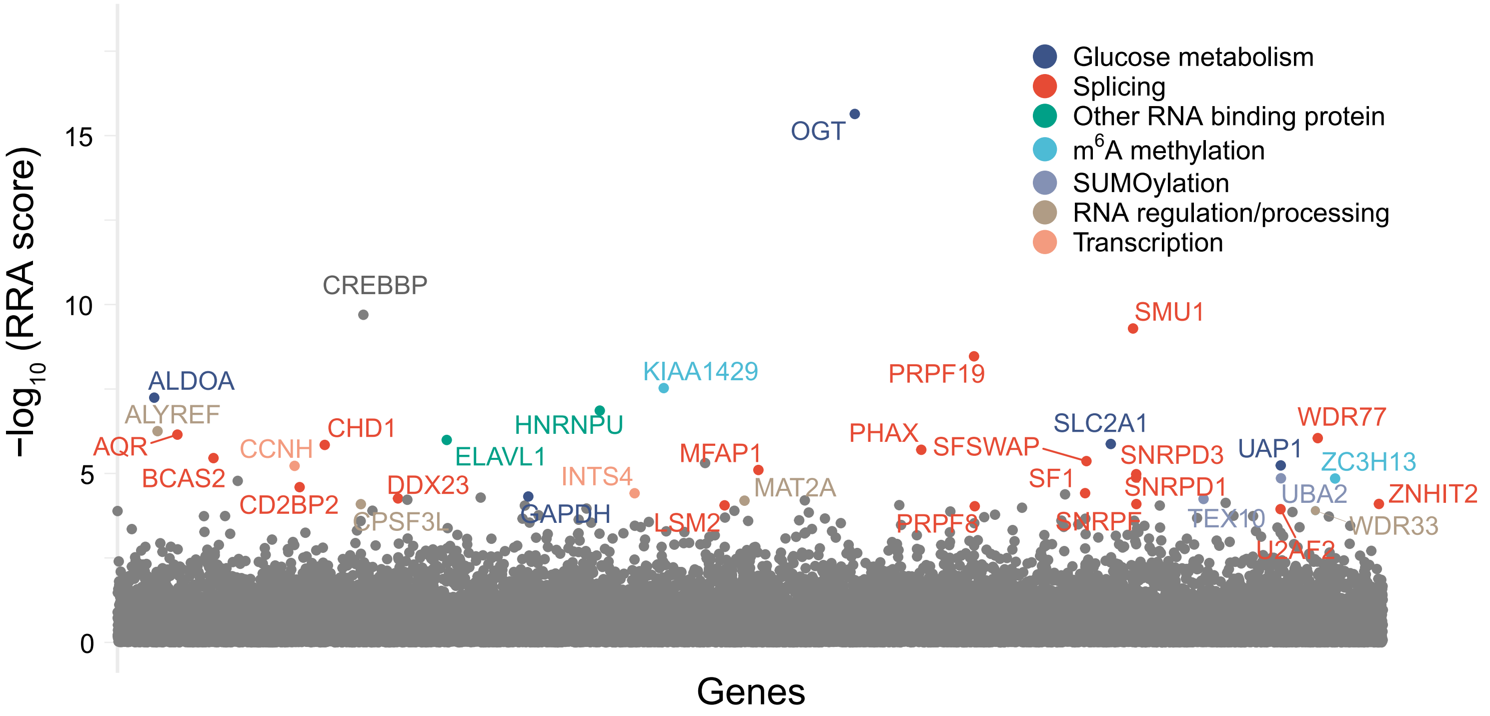
Glucose metabolism
Splicing
Other RNA binding protein
m6A methylation
SUMOylation
RNA regulation/processing
Transcription
OGT
CREBBP
SMU1
PRPF19
KIAA1429
ALDOA
ALYREF
AQR
CHD1
HNRNPU
ELAVL1
CCNH
BCAS2
CD2BP2
DDX23
INTS4
MFAP1
PHAX
SFSWAP
SLC2A1
SNRPD3
UAP1
WDR77
ZC3H13
SF1
SNRPD1
MAT2A
UBA2
ZNHIT2
GAPDH
CPSF3L
LSM2
PRPF8
SNRPF
TEX10
WDR33
U2AF2
15
10
5
0
−log10 (RRA score)
Genes

**Fig. S2**

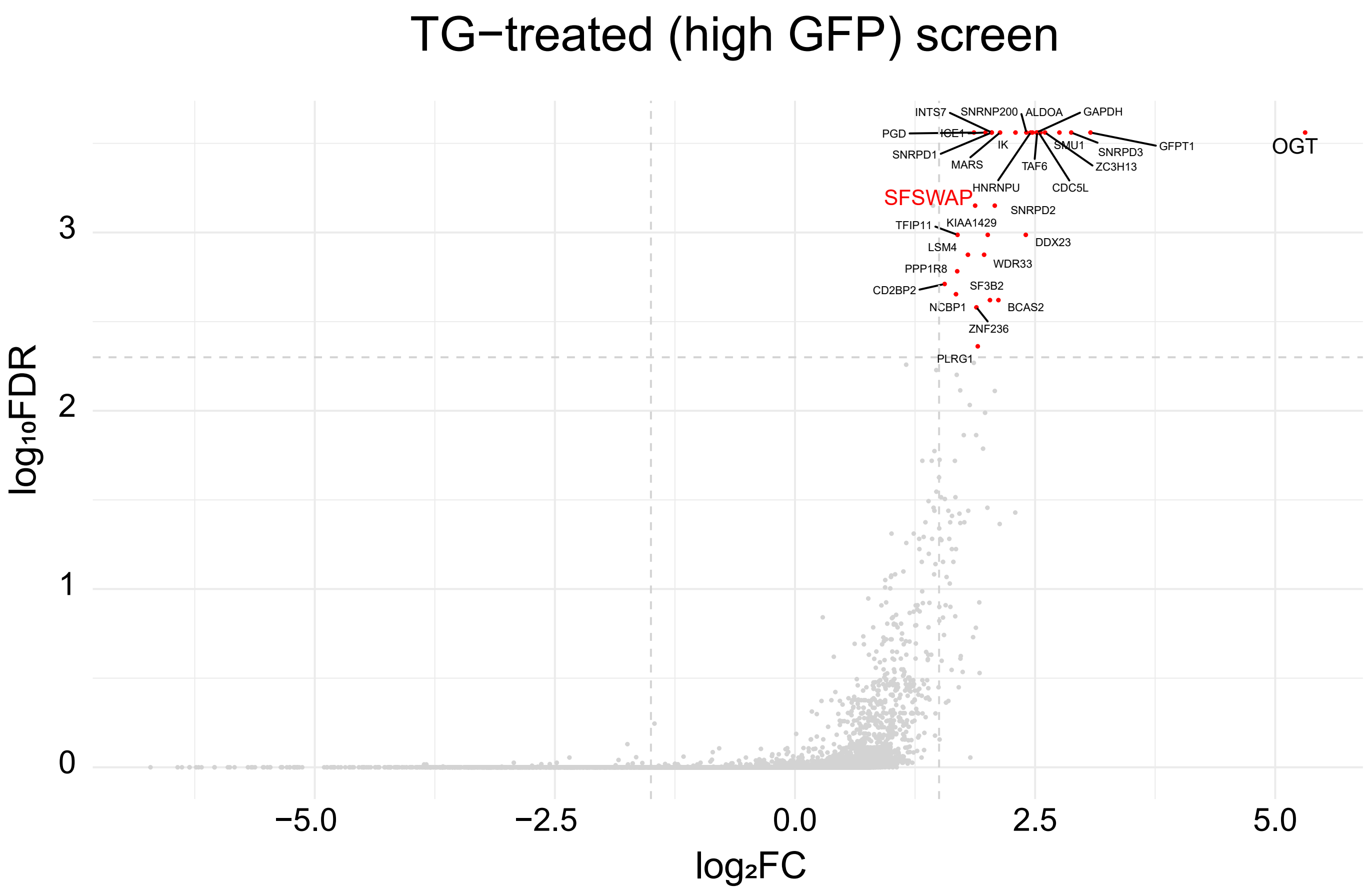
TG−treated (high GFP) screen
log₁₀FDR
log₂FC
OGT
SFSWAP
INTS7
SNRNP200
ALDOA
GAPDH
PGD
ICE1
IK
SMU1
SNRPD3
GFPT1
SNRPD1
MARS
TAF6
ZC3H13
HNRNPU
CDC5L
SNRPD2
TFIP11
KIAA1429
DDX23
LSM4
WDR33
PPP1R8
CD2BP2
SF3B2
NCBP1
BCAS2
ZNF236
PLRG1

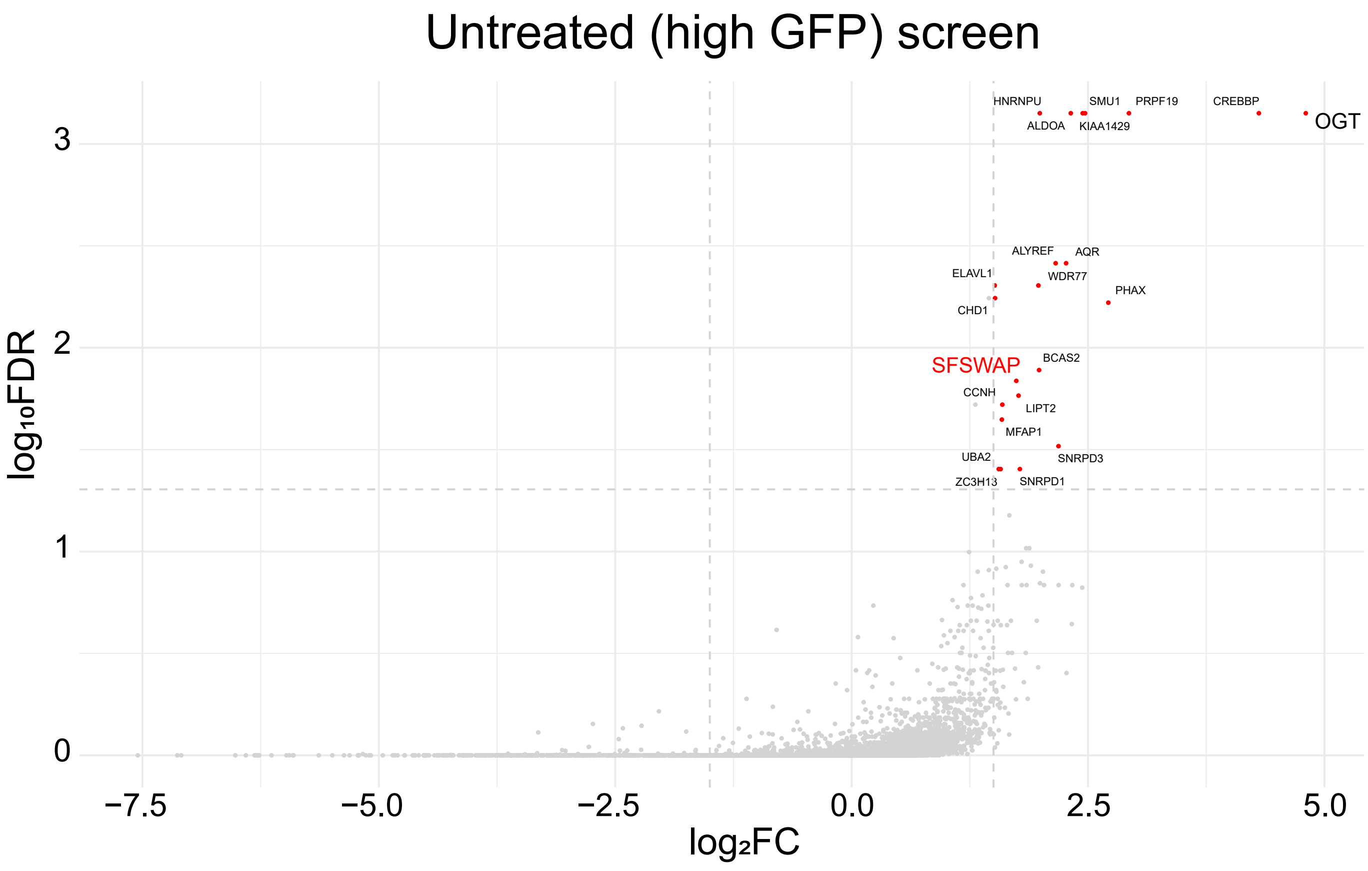
Untreated (high GFP) screen
log₁₀FDR
log₂FC
HNRNPU
SMU1
PRPF19
CREBBP
ALDOA
KIAA1429
OGT
ALYREF
AQR
ELAVL1
WDR77
PHAX
CHD1
SFSWAP
BCAS2
CCNH
LIPT2
MFAP1
UBA2
SNRPD3
ZC3H13
SNRPD1

**Fig. S3**

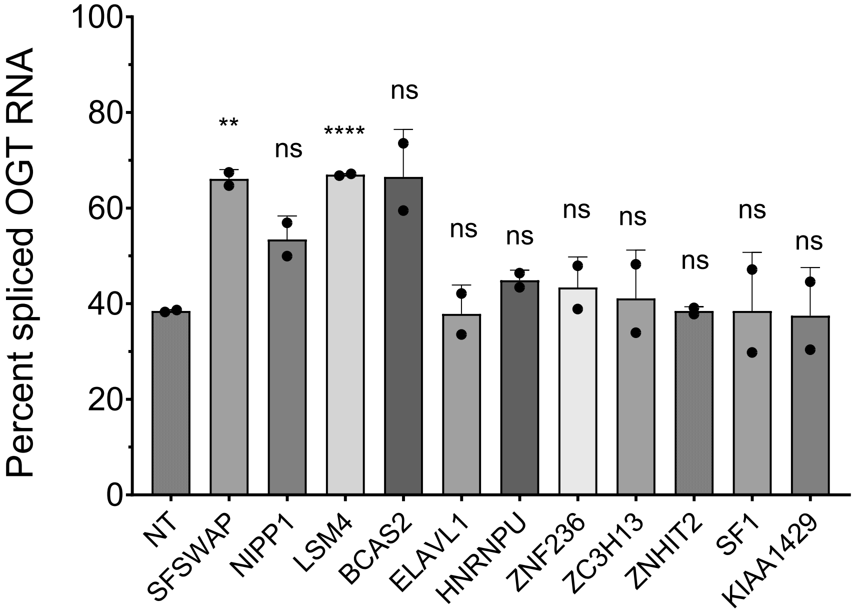

Percent spliced OGT RNA
100
80
60
40
20
0
**
ns
****
ns
ns
ns
ns
ns
ns
ns
ns
NT
SFSWAP
NIPP1
LSM4
BCAS2
ELAVL1
HNRNPU
ZNF236
ZC3H13
ZNHIT2
SF1
KIAA1429

**Fig. S4**

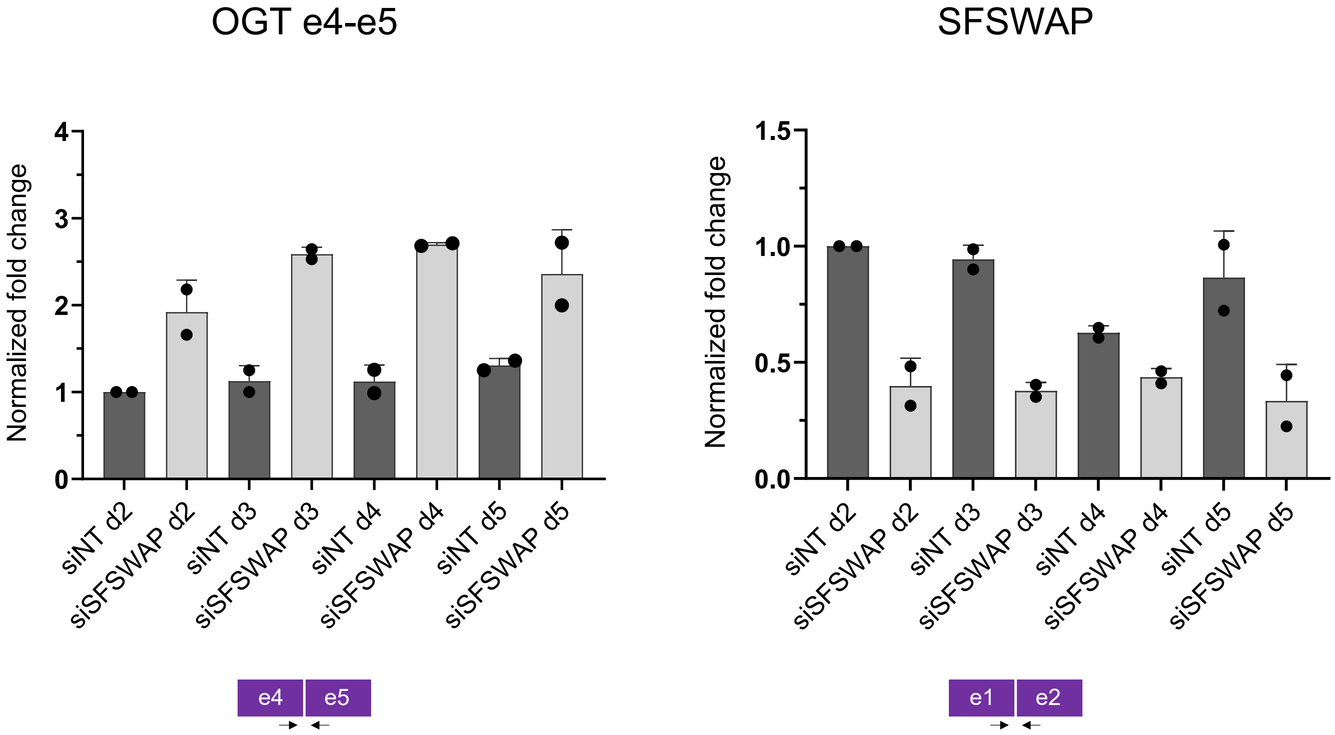

OGT e4-e5
Normalized fold change
0
1
2
3
4
siNT d2
siSFSWAP d2
siNT d3
siSFSWAP d3
siNT d4
siSFSWAP d4
siNT d5
siSFSWAP d5
e4
e5
SFSWAP
Normalized fold change
0.0
0.5
1.0
1.5
siNT d2
siSFSWAP d2
siNT d3
siSFSWAP d3
siNT d4
siSFSWAP d4
siNT d5
siSFSWAP d5
e1
e2

**Fig. S5**

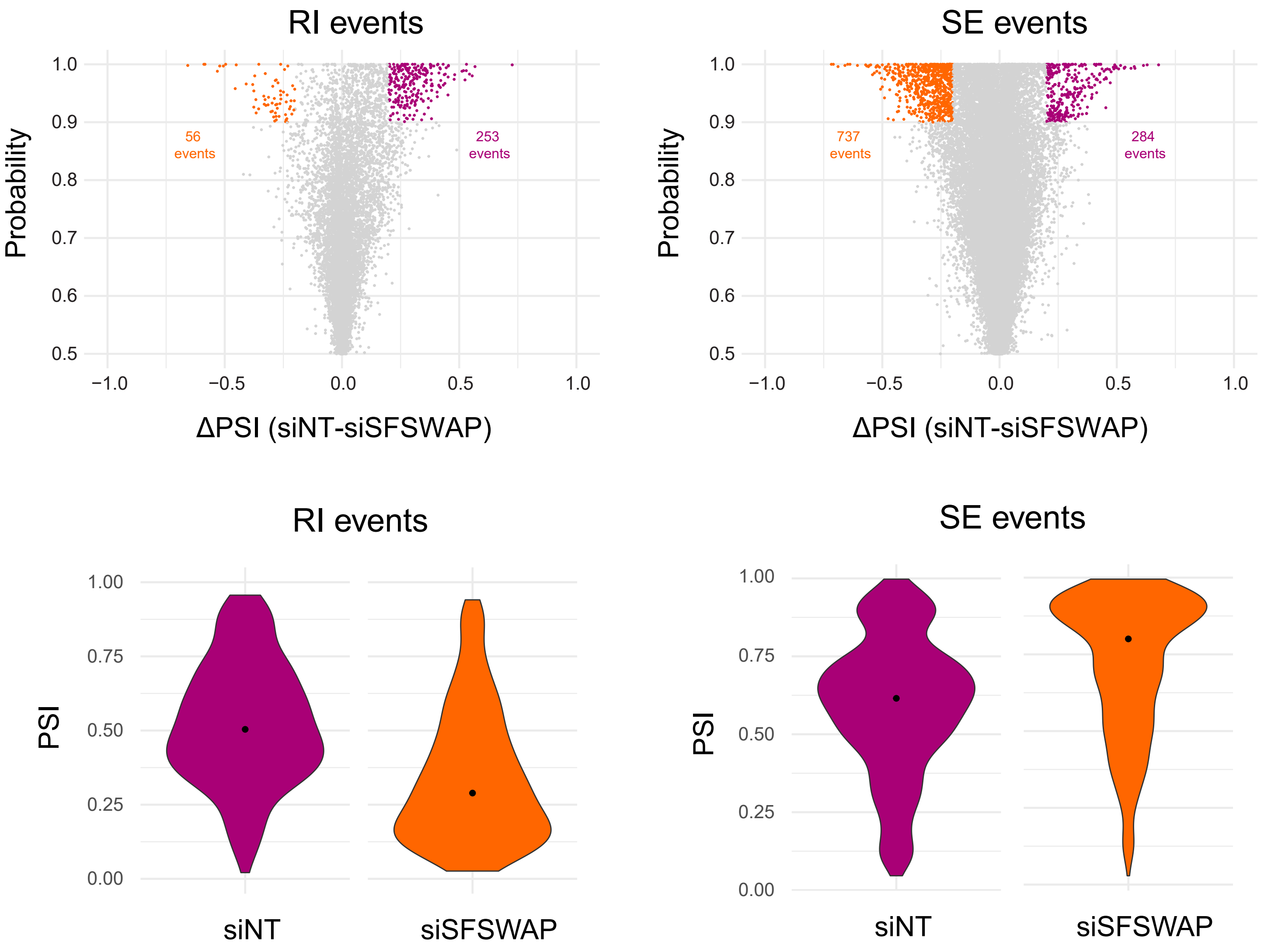

RI events
SE events
Probability
1.0
0.9
0.8
0.7
0.6
0.5
56 events
253 events
737 events
284 events
−1.0
−0.5
0.0
0.5
1.0
ΔPSI (siNT-siSFSWAP)
RI events
SE events
PSI
1.00
0.75
0.50
0.25
0.00
siNT
siSFSWAP

**Fig. S6**

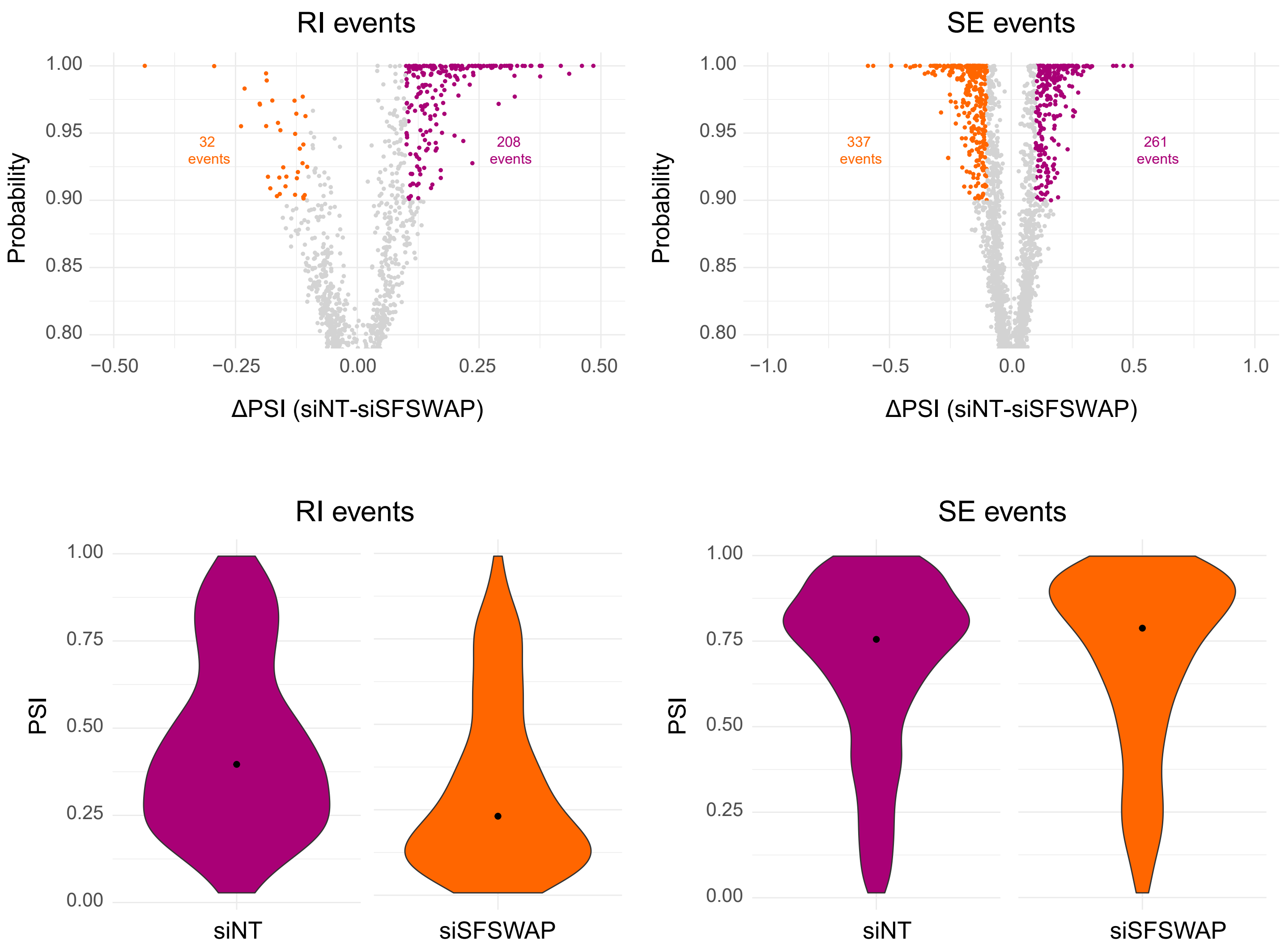

RI events
1.00
0.95
0.90
0.85
0.80
Probability
32
events
208
events
−0.50
−0.25
0.00
0.25
0.50
ΔPSI (siNT-siSFSWAP)
SE events
1.00
0.95
0.90
0.85
0.80
Probability
337
events
261
events
−1.0
−0.5
0.0
0.5
1.0
ΔPSI (siNT-siSFSWAP)
RI events
1.00
0.75
0.50
0.25
0.00
PSI
siNT
siSFSWAP
SE events
1.00
0.75
0.50
0.25
0.00
PSI
siNT
siSFSWAP

**Fig. S7**

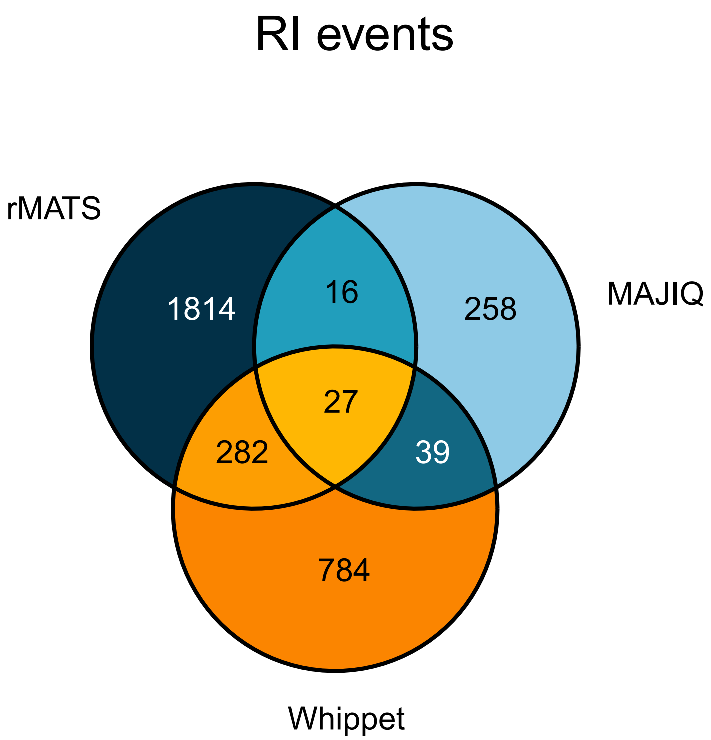

RI events
rMATS
MAJIQ
Whippet
1814
16
258
27
282
39
784
SE events
rMATS
MAJIQ
Whippet
6002
282
241
480
1307
128
3608

#### Fig. S8

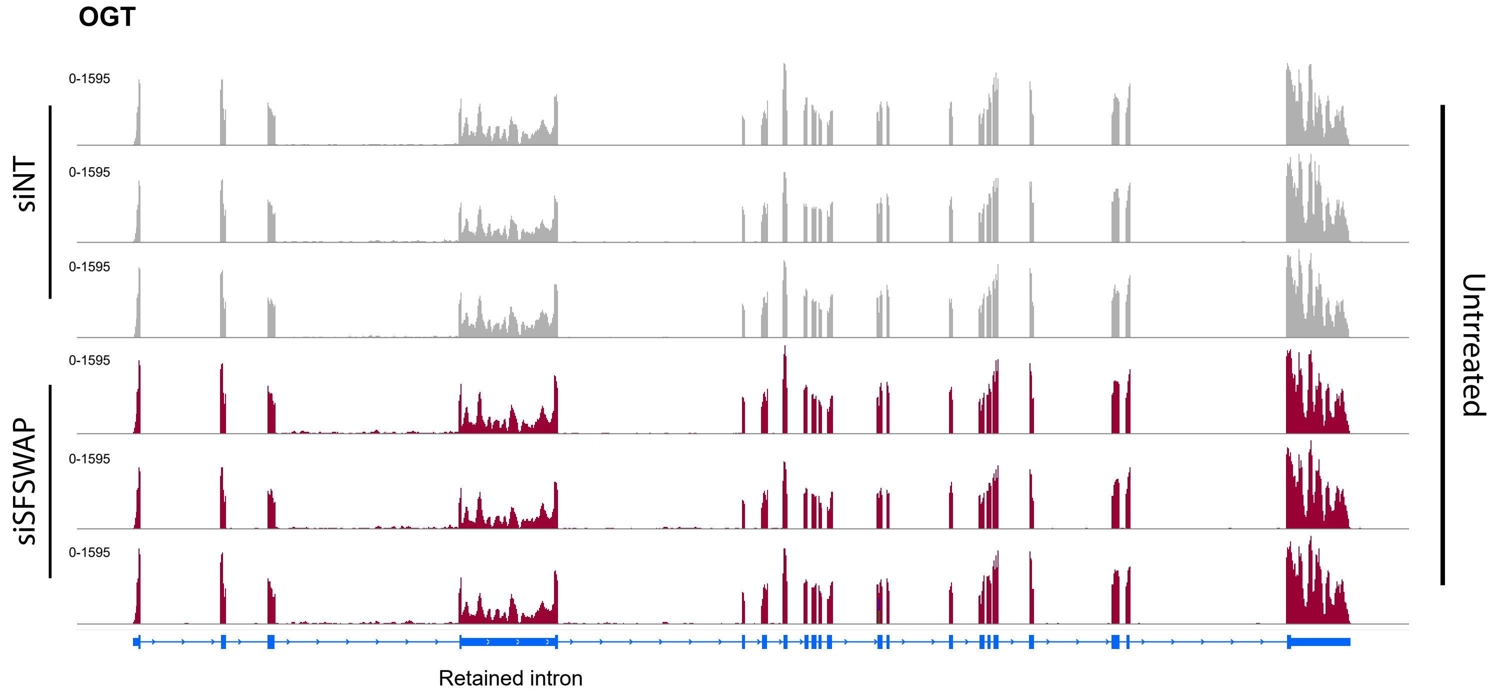
OGT
0-1595
0-1595
0-1595
0-1595
0-1595
0-1595
siNT
siSFSWAP
Untrreated
Retained intron

**Fig. S9**

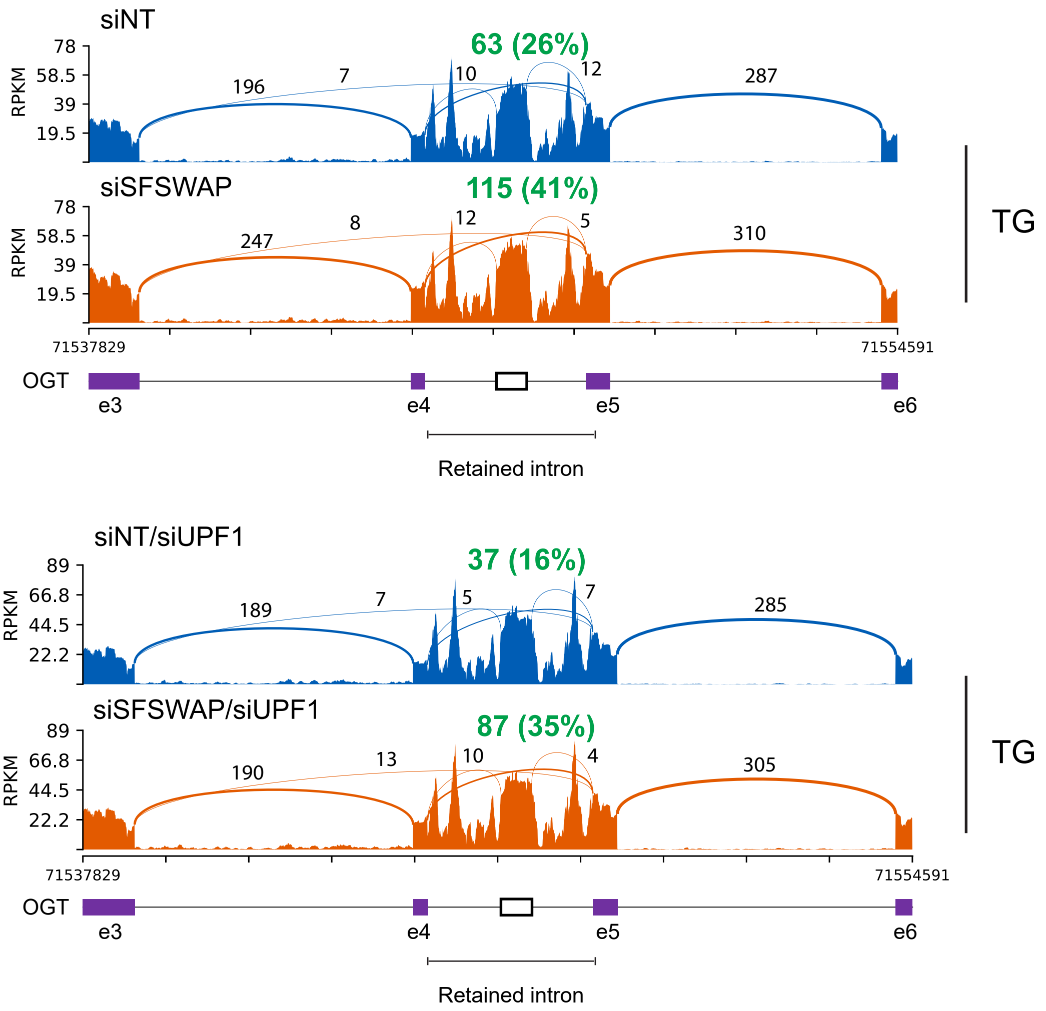
siNT
63 (26%)
RPKM
78
58.5
39
19.5
196
7
10
12
287
siSFSWAP
115 (41%)
247
8
12
5
310
TG
71537829
71554591
OGT
e3
e4
e5
e6
Retained intron
siNT/siUPF1
37 (16%)
89
66.8
44.5
22.2
189
7
5
7
285
siSFSWAP/siUPF1
87 (35%)
190
13
10
4
305
TG
71537829
71554591
OGT
e3
e4
e5
e6
Retained intron

**Fig. S10**

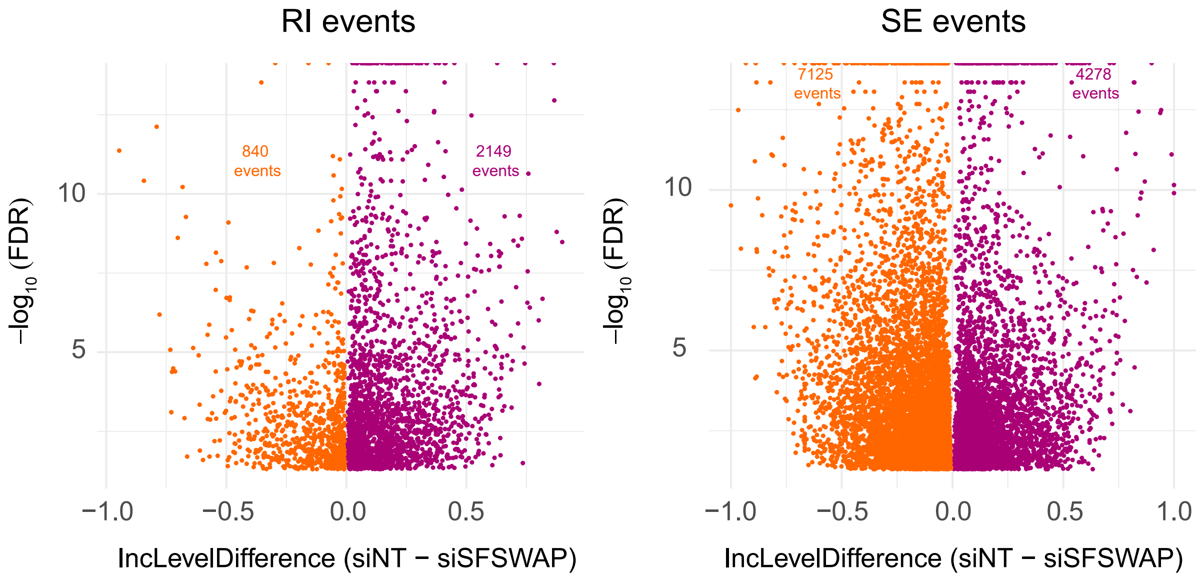
RI events
840
events
2149
events
−log10 (FDR)
10
5
−1.0
−0.5
0.0
0.5
IncLevelDifference (siNT − siSFSWAP)
SE events
7125
events
4278
events
−log10 (FDR)
10
5
−1.0
−0.5
0.0
0.5
1.0
IncLevelDifference (siNT − siSFSWAP)

**Fig. S11**

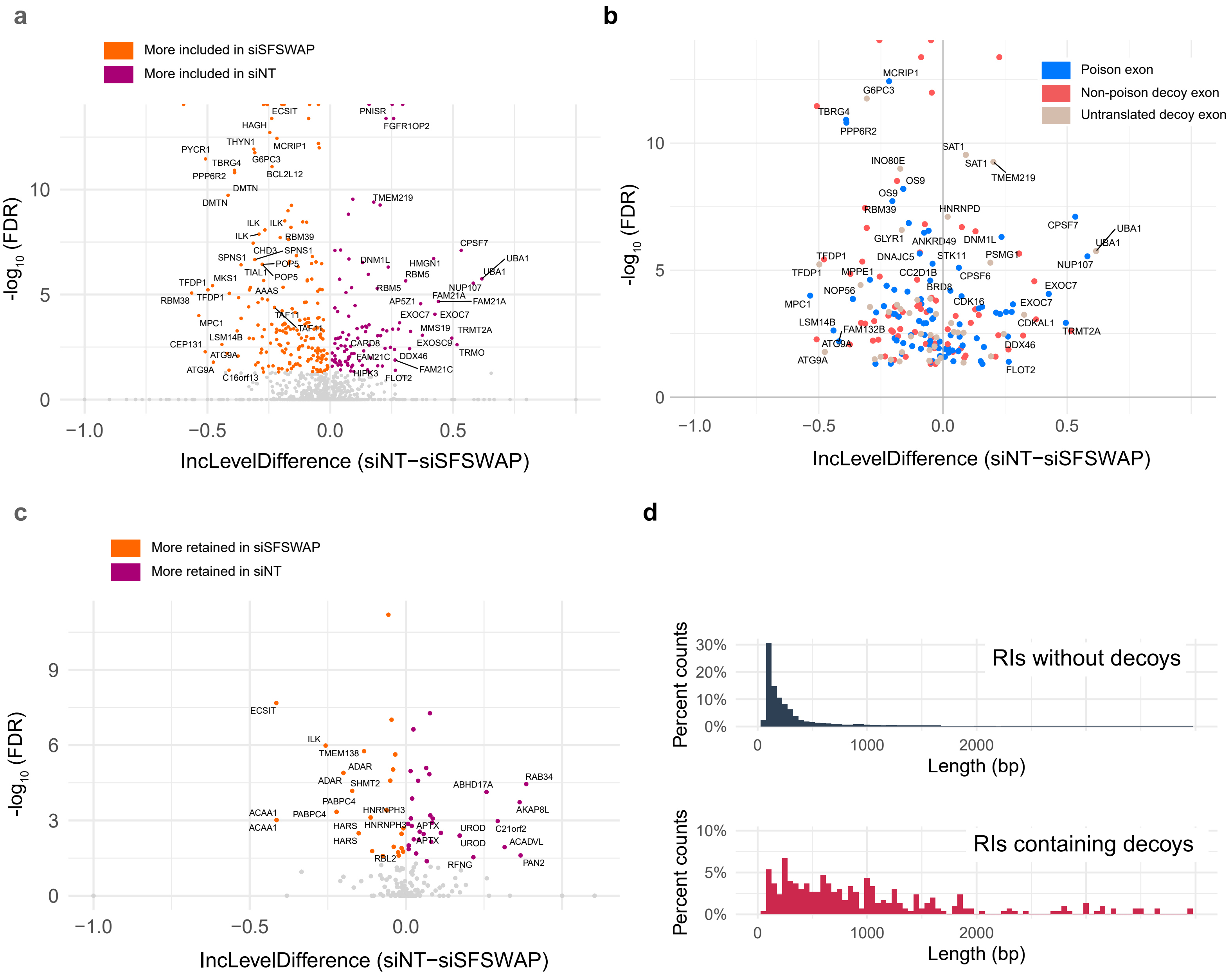
a
More included in siSFSWAP
More included in siNT
-log10 (FDR)
IncLevelDifference (siNT−siSFSWAP)
b
Poison exon
Non-poison decoy exon
Untranslated decoy exon
-log10 (FDR)
IncLevelDifference (siNT−siSFSWAP)
c
More retained in siSFSWAP
More retained in siNT
-log10 (FDR)
IncLevelDifference (siNT−siSFSWAP)
d
RIs without decoys
Percent counts
Length (bp)
RIs containing decoys
Percent counts
Length (bp)

### Fig. S12

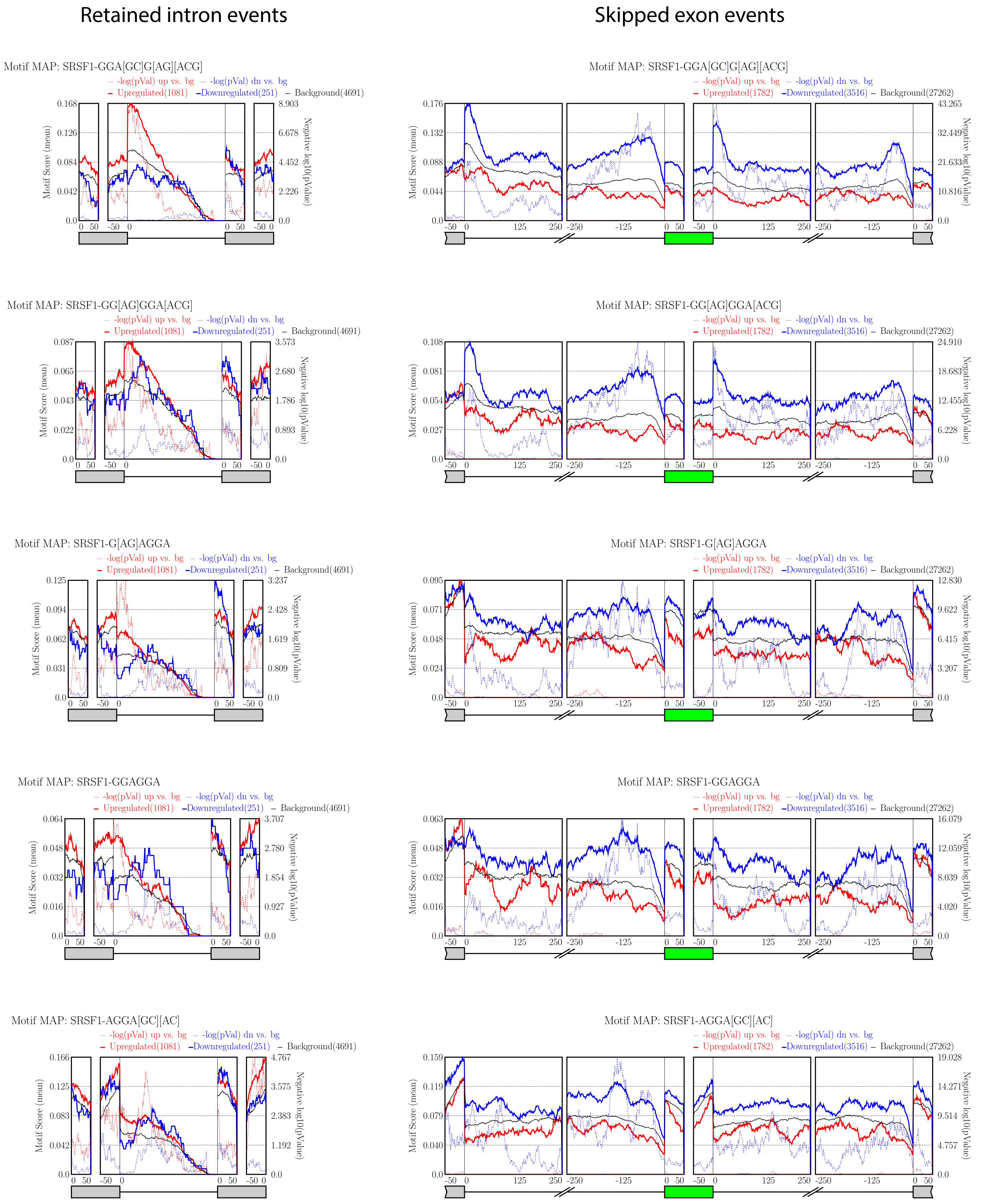

Retained intron events
Skipped exon events
Motif MAP: SRSF1-GGA[GC]G[AG][ACG]
-log(pVal) up vs. bg
-log(pVal) dn vs. bg
Upregulated(1081)
Downregulated(251)
Background(4691)
Upregulated(1782)
Downregulated(3516)
Background(27262)
Motif Score (mean)
Negative log10(pValue)
Motif MAP: SRSF1-GG[AG]GGA[ACG]
Motif MAP: SRSF1-G[AG]AGGA
Motif MAP: SRSF1-GGAGGA
Motif MAP: SRSF1-AGGA[GC][AC]

**Fig. S13**

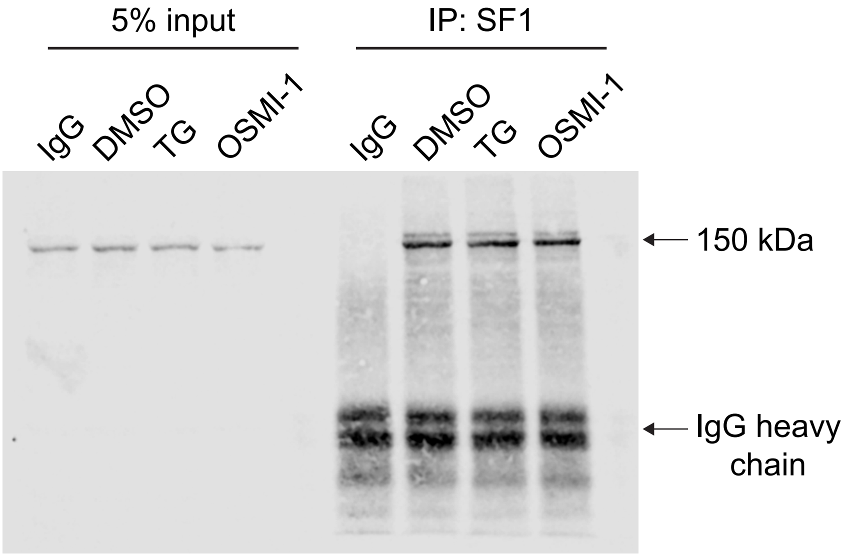


**α-SFSWAP**

**Fig. S14**

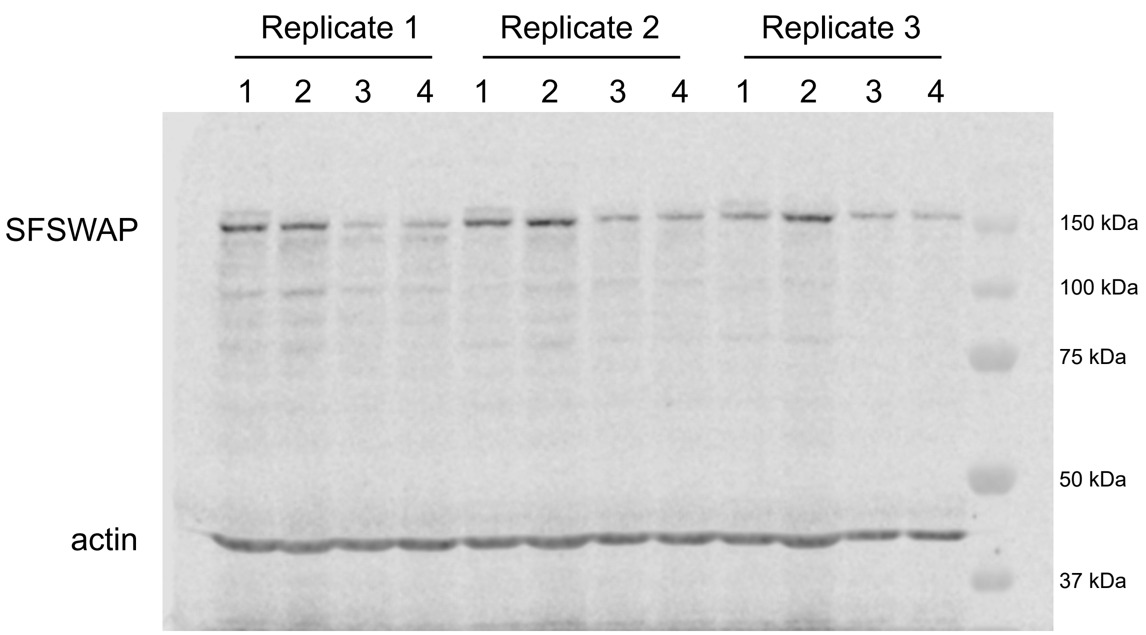


1. siNT
2. siSFSWAP (siRNA #1)
3. siSFSWAP (siRNA #2)
4. siSFSWAP (siRNA #1+2) (used for RNA-seq experiments)

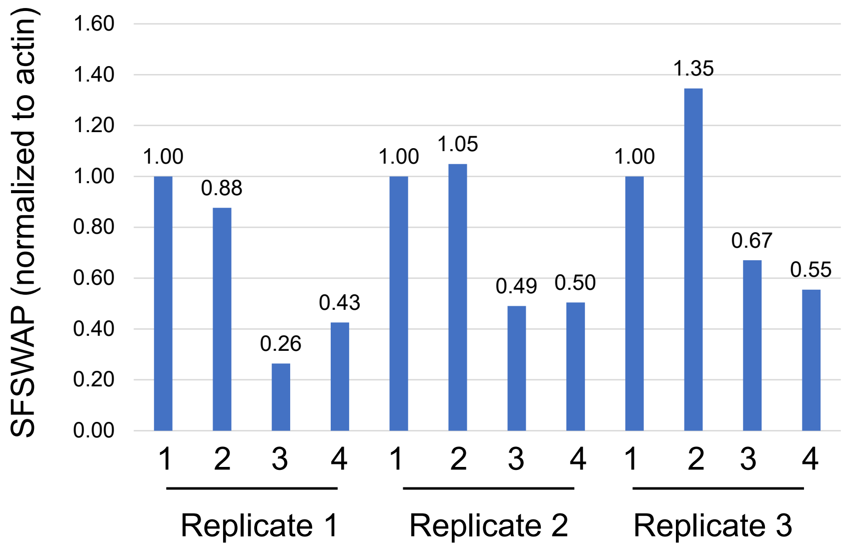

#### Fig. S15

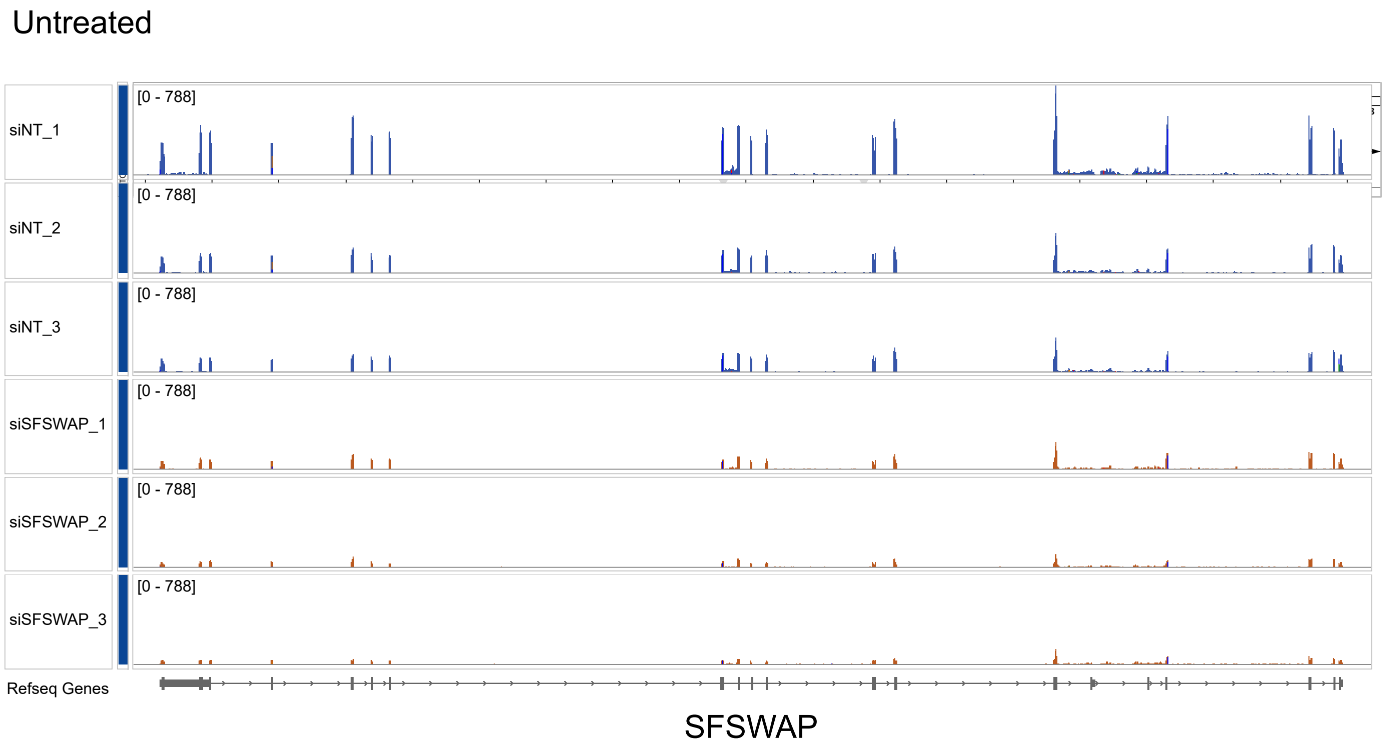
Untreated
siNT_1
[0 - 788]
siNT_2
[0 - 788]
siNT_3
[0 - 788]
siSFSWAP_1
[0 - 788]
siSFSWAP_2
[0 - 788]
siSFSWAP_3
[0 - 788]
Refseq Genes
SFSWAP

Fig. S16

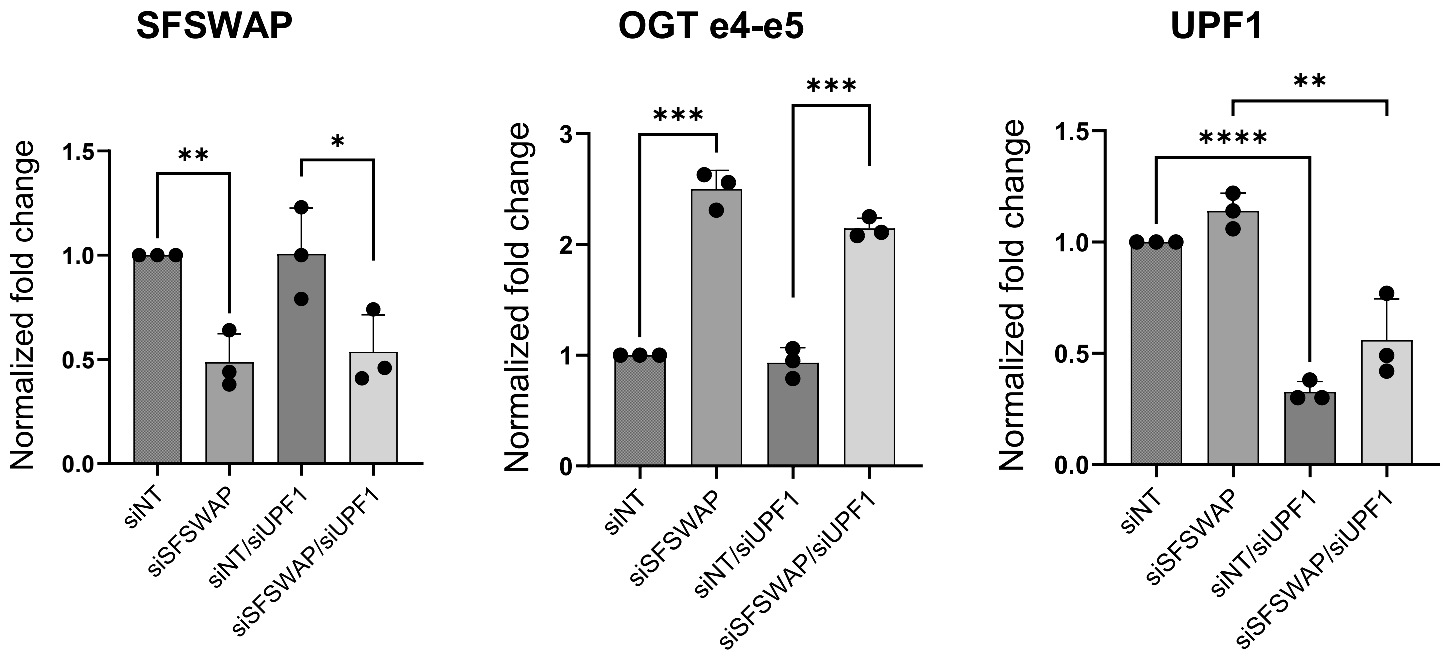

SFSWAP
OGT e4-e5
UPF1
Normalized fold change
**
*
***
***
****
**
siNT
siSFSWAP
siNT/siUPF1
siSFSWAP/siUPF1
0.0
0.5
1.0
1.5
0
1
2
3

**Fig. S17**

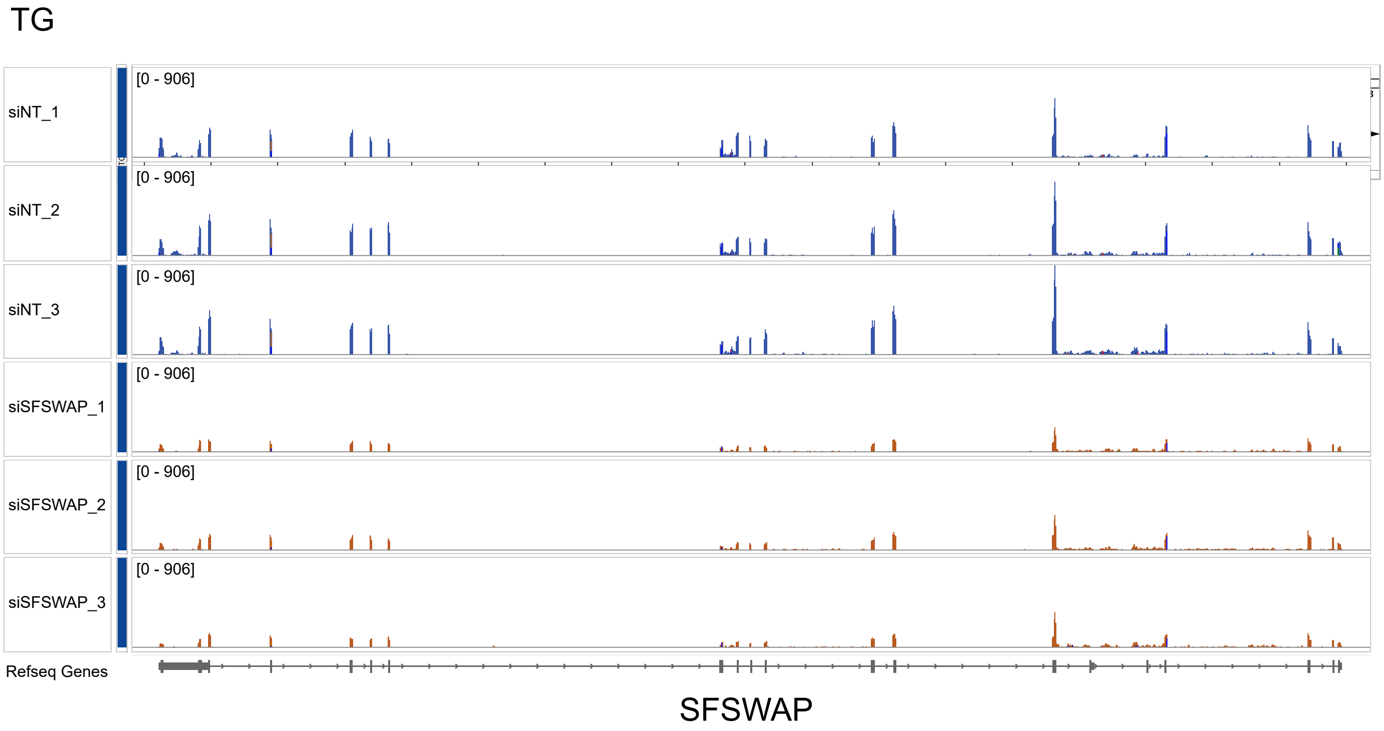

TG
siNT_1
siNT_2
siNT_3
siSFSWAP_1
siSFSWAP_2
siSFSWAP_3
[0 - 906]
Refseq Genes
SFSWAP


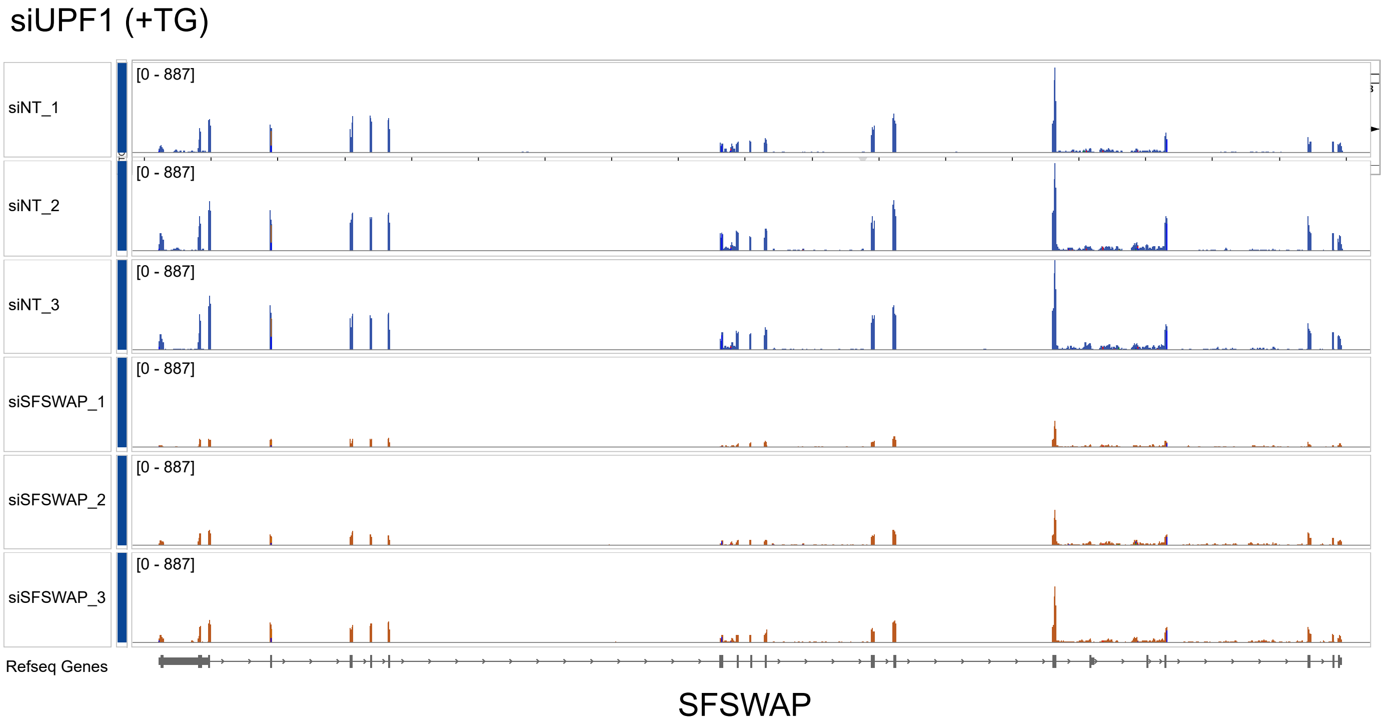

siUPF1 (+TG)
siNT_1
siNT_2
siNT_3
siSFSWAP_1
siSFSWAP_2
siSFSWAP_3
[0 - 887]
Refseq Genes
SFSWAP
